# scTREND: An annotation-free single-cell time-resolved and condition-dependent hazard model

**DOI:** 10.64898/2026.01.26.701686

**Authors:** Shintaro Yuki, Chikara Mizukoshi, Ko Abe, Teppei Shimamura

## Abstract

Prognosis in cancer and other complex diseases is shaped by heterogeneous cell states and clinical or genetic contexts whose effects change over time. Yet most survival analyses assume predefined cell types and time-invariant covariates, and cohorts pairing single-cell transcriptomes with outcomes are limited, obscuring within-type heterogeneity, time-varying risk, and condition-specific effects. We developed scTREND, an annotation-free, time-resolved, condition-dependent hazard model that learns variational single-cell embeddings, deconvolves bulk or spatial samples without labels, and fits a conditional piecewise-constant hazard model to estimate cell-level hazard coefficients across discrete time bins and conditions. scTREND enables cell-level risk attribution without prior cell-type annotations, dynamic risk modeling over time, and mutation- or treatment-specific risk assessment. In simulations, scTREND recovered ground-truth temporal and condition-specific coefficients with strong concordance and improved prediction over an existing method. In melanoma, scTREND identified subpopulations with risk amplified in BRAF-mutant tumors; in COVID-19, cytotoxic T-cell subpopulations with early-to-late risk reversal and pathways associated with prognosis under remdesivir; and in spatial transcriptomics of clear cell renal carcinoma, spatial regions whose prognostic relevance shifts over time. Across diseases and modalities (bulk RNA-seq and spatial transcriptomics), scTREND provides a time-resolved, condition-aware link between single-cell states and clinical outcomes and is implemented in Python (GitHub:https://github.com/R301Carbine/scTREND).

## Introduction

Recent studies have demonstrated that the prognosis of cancer patients is profoundly influenced by the diversity of cellular populations within tumors and by the composition of the tumor microenvironment [1]. Indeed, immunological indicators such as the type and density of infiltrating immune cells have been reported to serve as powerful prognostic factors [2]. These observations highlight the importance of accurately characterizing the heterogeneity of tumor-associated cells in order to predict patient survival outcomes. Advances in single-cell RNA sequencing (scRNA-seq) technology have enabled comprehensive profiling of the gene expression programs of individual cells within tumors. As a result, it has become evident that heterogeneity in both malignant and immune cells influences patient prognosis and therapeutic responses. For example, scRNA-seq studies have identified specific immune cell subgroups associated with response to immune checkpoint blockade therapy as well as with poor clinical outcomes [3], highlighting the promise of approaches that identify prognostically relevant cellular populations at single-cell resolution. Furthermore, studies in breast and other cancers have shown that prognostic factors can be time-dependent [4] and that risk contributions may vary according to driver gene mutations or prior treatment history [5]. Taken together, these findings suggest that the biological heterogeneity within tumors and its temporal and condition-dependent dynamics play a critical role in shaping patient outcomes. A comprehensive understanding of such heterogeneity is therefore essential for elucidating the mechanisms underlying cancer prognosis.

Despite these advances, large-scale cohorts that integrate scRNA-seq data with clinical outcome information remain scarce due to technical and financial limitations. By contrast, bulk RNA-seq datasets linked to clinical outcomes have been systematically generated from large patient cohorts, most notably The Cancer Genome Atlas (TCGA), and are widely utilized for prognostic analyses. In this context, numerous deconvolution methods have been developed to infer cell-type proportions from bulk RNA-seq data, enabling investigations into the associations between immune cell infiltration or cellular composition and prognosis [6, 7]. However, most of these approaches rely on predefined cell-type categories and are therefore limited in their ability to capture cellular heterogeneity within a given type. Consequently, subtle differences in cellular state or the contributions of specific subclusters to prognosis may be overlooked. Moreover, survival analysis in these studies typically employs the Cox proportional hazards model, which assumes that covariate effects remain constant over time and across patient conditions. To relax these constraints, several neural network-based survival models such as DeepSurv [8] and Cox-nnet [9] have been proposed, which allow flexible modeling of nonlinear relationships between covariates and risk. Even with this flexibility, these methods cannot capture temporal and condition-dependent effects described above, potentially obscuring clinically important cellular populations or molecular pathways that influence prognosis.

To address these challenges, we developed scTREND, a novel framework for modeling time-resolved and condition-dependent hazards at single-cell resolution. Compared to conventional survival analysis models, our framework possesses the following three distinct advantages.

First, scTREND enables single-cell resolution analysis without requiring cell-type annotations. Most conventional methods rely on analyses based on predefined cell types, which limits their ability to capture heterogeneity within cell types or the prognostic contributions of specific subclusters. In contrast, the proposed method employs a variational autoencoder (VAE) to learn latent representations of cells, allowing estimation of risk contributions at the individual cell level without dependence on prior cell-type annotations. This enables the discovery of fine-grained cell populations associated with prognosis, unconstrained by broad cell-type categories.

Second, scTREND enables dynamic modeling of risk changes over time. The conventional Cox proportional hazards model assumes that the effects of covariates remain constant over time. Furthermore, while neural network-based models such as DeepSurv can handle nonlinear relationships between covariates and risk, they cannot capture time-dependent effects. By adopting a conditional piecewise-constant hazard model, our approach enables estimation of cell-level risk contributions that change over time. This makes it possible to identify cell populations and pathways whose prognostic impact differs between early and late stages of disease.

Third, scTREND enables risk assessment dependent on patient-specific conditions by modeling prognostic effects—such as those driven by driver gene mutations or therapeutic interventions—that conventional models fail to capture. By incorporating these conditions into the model, it becomes possible to identify cell populations and molecular pathways whose prognostic contributions are amplified or diminished only under specific conditions.

Furthermore, scTREND can extend this framework to spatial transcriptomics data (such as Visium), enabling the identification of spatial regions whose prognostic contributions change over time, which represents another significant advantage.

Through numerical simulations, scTREND demonstrated the ability to accurately reconstruct the underlying structure of the true hazard ratio. When applied to melanoma data, it identified cell populations whose prognostic risks were boosted by mutations in the driver gene, as well as those exhibiting temporal variations in risk contributions. Furthermore, analysis of COVID-19 datasets revealed cell populations whose prognostic risks were altered by the administration of the antiviral drug remdesivir, along with the molecular features associated with these cells. Finally, in the application to Visium data from renal cell carcinoma, scTREND identified spatial regions whose contributions to prognosis vary over time, as well as the genes that drive these prognostic dynamics. This framework enables the identification of specific cells with time- and condition-dependent contributions to prognosis, as well as the quantitative evaluation of variations in their contribution levels. As a result, scTREND reveals cellular populations and gene signatures with strong prognostic relevance, thereby offering novel opportunities for the discovery of therapeutic targets.

## Results

### Overview of scTREND

We developed scTREND, a novel deep generative framework (Figure 1). This framework first learns low-dimensional representations of cellular states from scRNA-seq data and then projects these learned states onto patient samples, enabling the quantification of cellular composition in bulk RNA-seq cohorts. By integrating these estimates with a time-resolved survival model, scTREND produces patient-specific risk scores and identifies prognosis-associated cellular programs that may vary across conditions and time intervals.

**Fig. 1.**
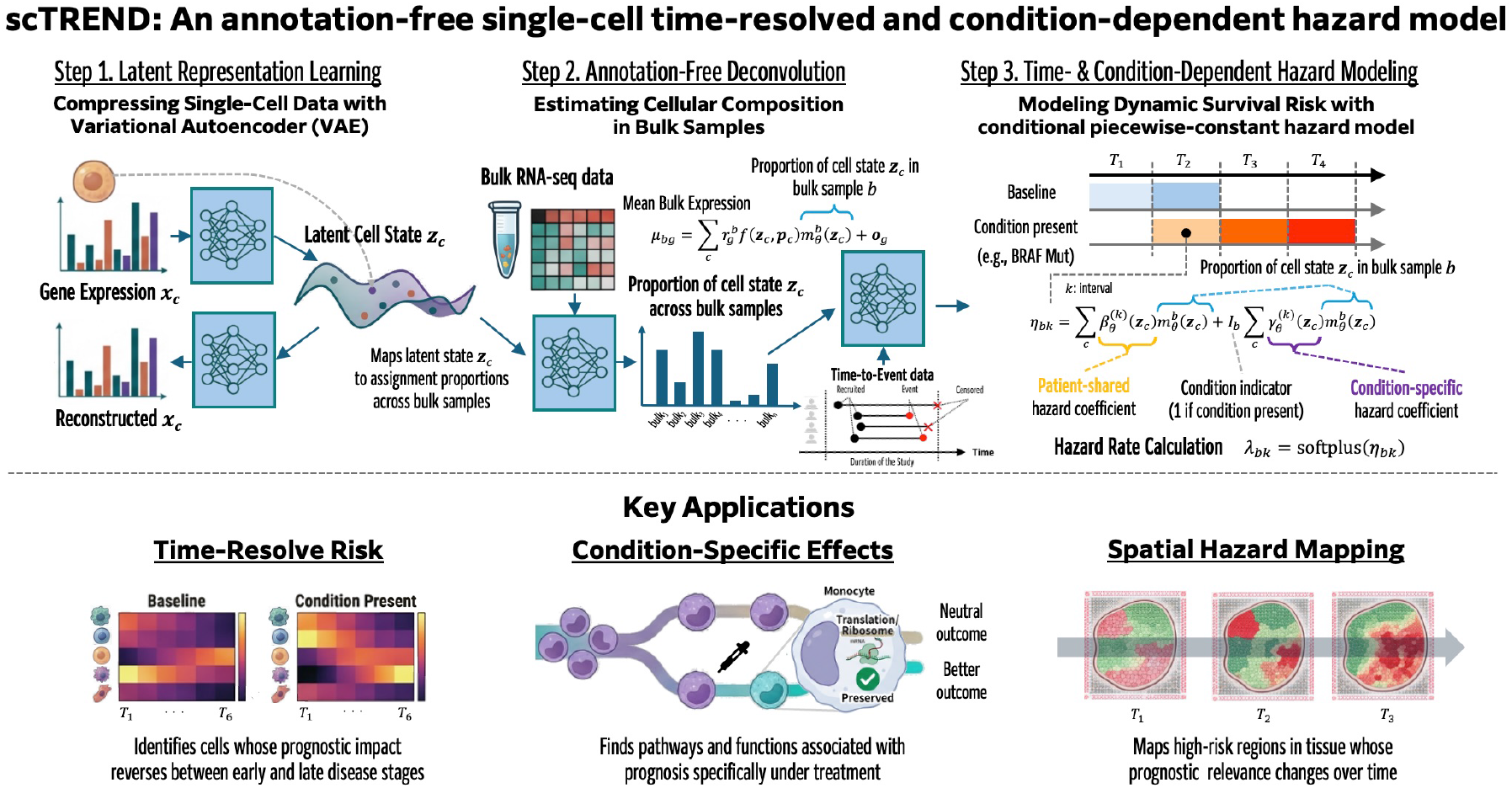
Overview of scTREND. Conceptual framework of scTREND. Our framework estimates the contributions of individual cells to patient prognosis across different conditions and time points. It leverages VAE-derived single-cell latent representations together with bulk-level cell-type mixing ratios and cell-level hazard coefficients that quantify each cell’s contribution to patient prognosis. This enables the computation of patient-level risk scores and the identification of cells whose prognostic impact varies over time and condition for each patient. Key applications include time-resolved identification of prognostic cell populations, condition-specific functional characterization of risk-associated pathways, and spatial hazard mapping of tissue regions with time-dependent clinical relevance.

Moreover, scTREND is readily applicable to spatial transcriptomics data. By assigning learned cellular states to individual spots, it can estimate spot-level risks and highlight spatial regions associated with prognosis over time, providing a unified approach for analyzing bulk and spatial transcriptomic cohorts within the same conceptual framework.

### Performance evaluation of scTREND on simulated data

#### Data generation and evaluation metrics

We evaluated the performance of scTREND using simulated data (Figure 2a). This model is designed to estimate hazard coefficients that vary over time while accounting for condition-dependent factors such as mutations or treatments. To this end, we defined cell clusters and specified the true condition- and time-dependent regression coefficients for each time interval.

**Fig. 2.**
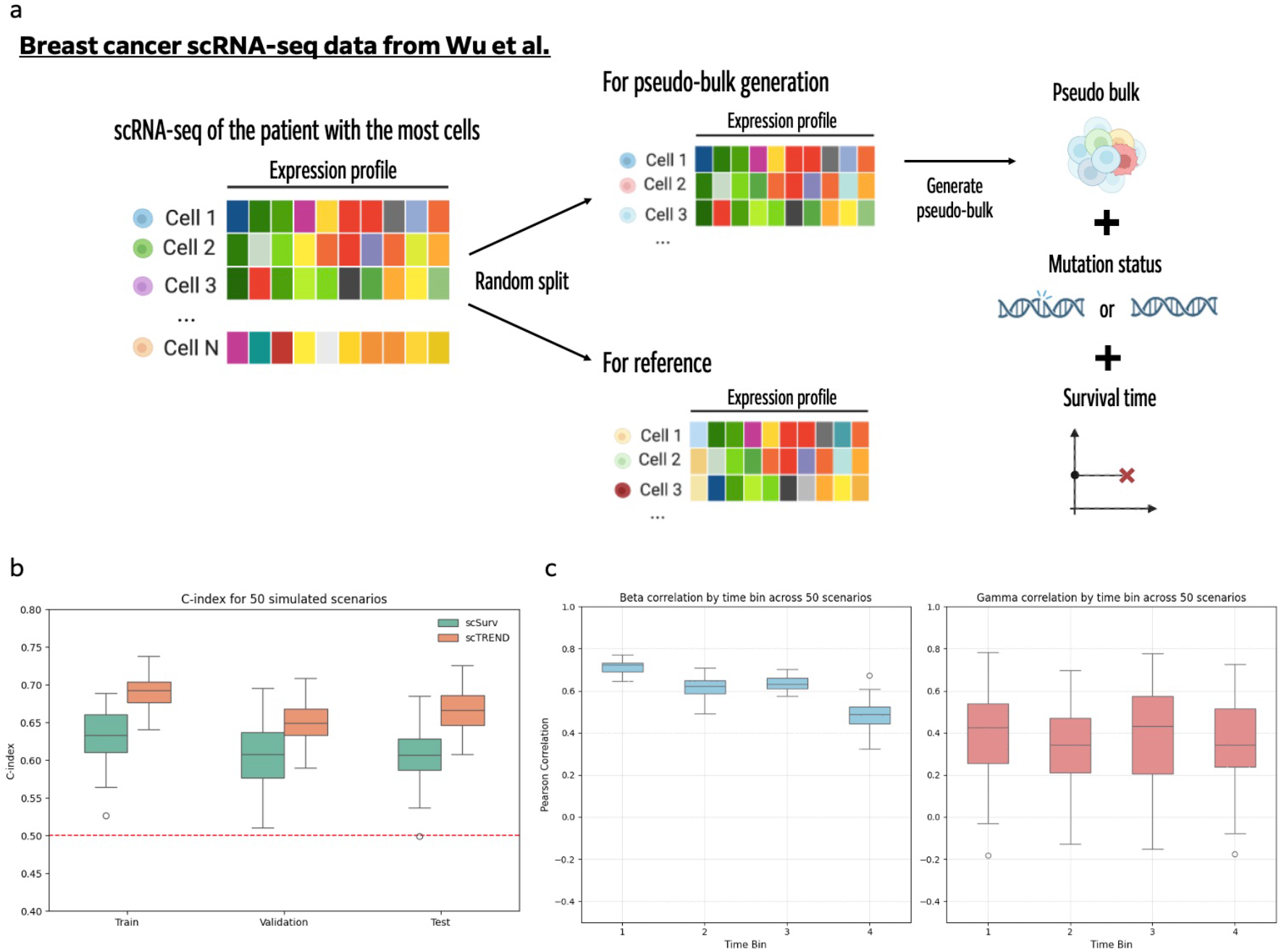
Performance evaluation of scTREND. (a) Data and splitting strategy. The breast cancer scRNA-seq dataset from [10] was used. Data from the patient with the largest number of cells were randomly divided into two subsets, one for pseudo-bulk generation and the other for reference. (b) Comparison with existing methods based on the c-index. Across 50 simulated scenarios, the performance of scTREND was compared with that of the existing method, scSurv. (c) Estimation accuracy of regression coefficients. For each of the 50 simulated scenarios, the Pearson correlation coefficients between the true and estimated regression coefficients were calculated for each time interval and visualized as boxplots. The left panel shows the results for 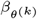, and the right panel shows those for 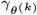.

Simulated data were generated using the breast cancer scRNA-seq dataset from [10], following the simulation procedure described in scSurv [11]. A detailed description of the data-generation procedure and evaluation metrics is provided in Methods.

Using the generated pseudo-bulk samples, we varied the true regression coefficients and simulated 50 distinct sets of survival times. To evaluate the estimation performance, we computed the Pearson correlation between the true and estimated regression coefficients, as well as the concordance index (c-index) based on the risk score 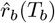.

#### Evaluation results

To validate the predictive performance and estimation accuracy of scTREND, we conducted a comparative evaluation with scSurv, an existing single-cell-level survival modeling method. The c-index values confirmed that our method achieved robust predictive performance on the test data and consistently outperformed scSurv (Figure 2b). This result demonstrates that its time-resolved and condition-aware hazard modeling enables more accurate risk prediction compared to conventional single-cell survival analysis methods.

Furthermore, evaluation of the estimation accuracy of scTREND’s time-dependent and condition-dependent hazard coefficients revealed that the common regression coefficients 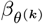 exhibited strong positive correlations with the ground truth across all time intervals (Figure 2c, left panel). Notably, the condition-dependent coefficients 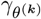 also maintained positive correlations in most scenarios (Figure 2c, right panel). This demonstrates that our method can appropriately separate and estimate patient-shared effects and condition-specific effects. Estimating such coefficients that differ across time intervals and conditions is fundamentally impossible with conventional Cox proportional hazards models or DeepSurv, and these results numerically substantiate the unique strengths of scTREND. Collectively, these results demonstrate the superior accuracy and robustness of our method in modeling survival risk from three perspectives: (1) predictive performance surpassing existing methods, (2) high-precision estimation of time-dependent coefficients, and (3) appropriate separation and estimation of condition-dependent coefficients. This indicates that it has a solid theoretical foundation in simulation data, and reliable estimation can be expected in applications to real datasets where risk changes dynamically with time or clinical context.

### Analysis of the impact of BRAF mutations in melanoma using scTREND

We applied scTREND to the melanoma cohort from TCGA. Melanoma is a malignant tumor originating from pigment-producing melanocytes in the skin and is characterized by high metastatic potential and substantial molecular heterogeneity [12]. Because it exhibits a high functional mutation burden primarily involving BRAF [13], we focused on BRAF mutations in this application. In this cohort, approximately 49.6% of bulk samples contained BRAF mutations. In addition, it has been reported that melanoma prognosis can evolve over follow-up, as conditional survival improves with increasing time since diagnosis [14]. These characteristics of the dataset provided an ideal opportunity to validate scTREND’s time-resolved and condition-aware modeling capabilities.

First, we used the scRNA-seq dataset of melanoma from [15] as a reference. The cell type annotations were adopted from the original study (Figure 3a). As a result of applying scTREND, the mean c-index across the training, validation, and test datasets was approximately 0.7 (Figure 3b). The estimation was performed with four predefined temporal intervals. The details of the time point settings are provided in the Methods section. The UMAP visualizations of the estimated 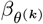 (***z***_*c*_) and 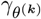 (***z***_*c*_) values at time bins 1-4 are provided in Supplementary Figures S1b and S1c, respectively.

**Fig. 3.**
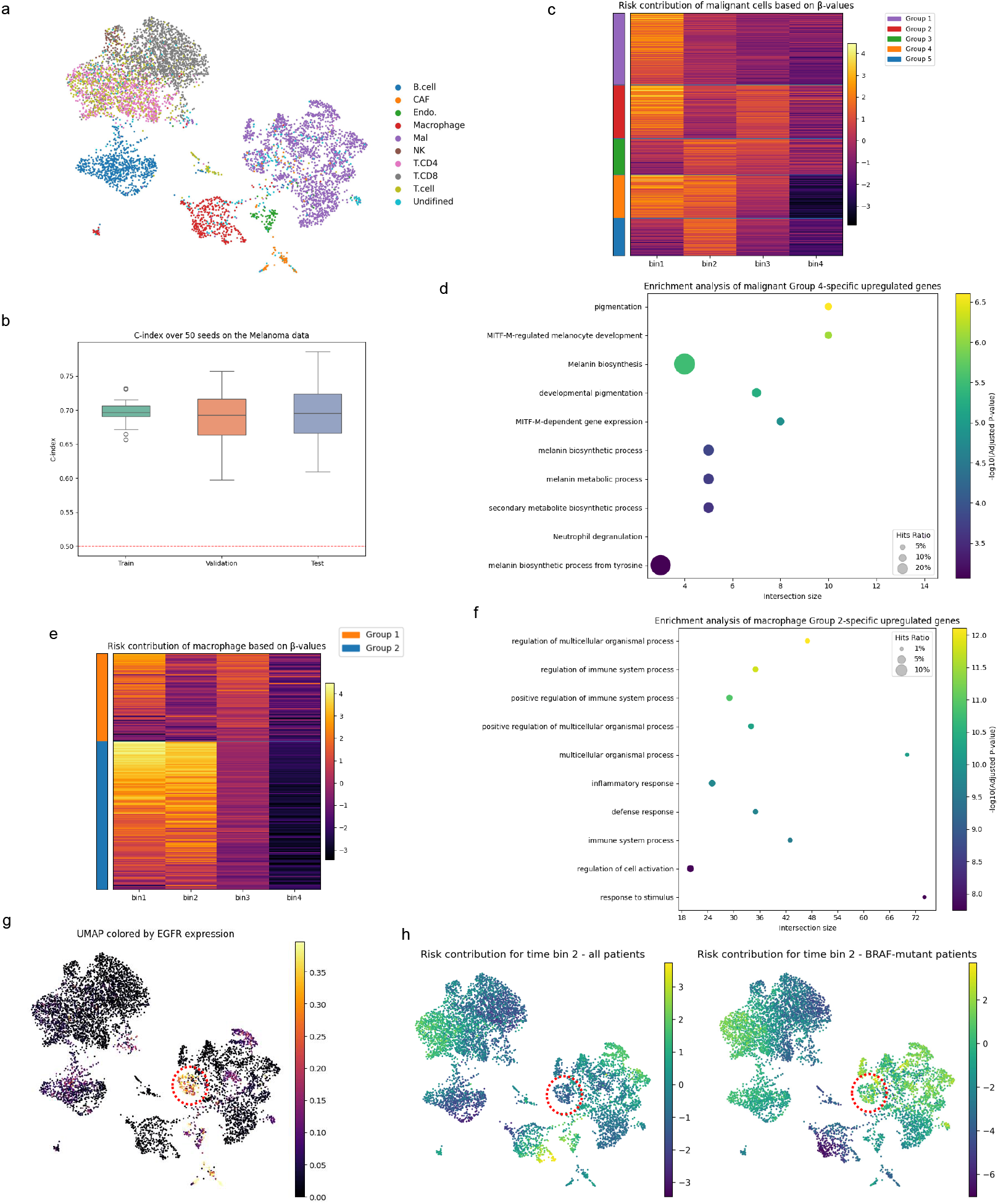
BRAF mutation-associated analysis in melanoma using scTREND. (a) UMAP visualization of cell types in the reference scRNA-seq dataset. (b) Boxplots of c-index values for the training, validation, and test datasets of the bulk data across 50 random splits. (c) Heatmap of 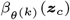 values for malignant cells across time bins. Clusters were identified on the basis of the estimated 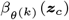 values. (d) Dot plot of enrichment analysis for the top 100 genes specifically upregulated in malignant cell Group 4 relative to other groups. Each dot represents an enriched gene set; the y-axis lists enriched terms, and the x-axis indicates the number of overlapping genes. Dot size reflects the ratio of overlapping genes to gene set size, and color intensity represents statistical significance. (e) Heatmap of 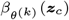 values for macrophage cells across time bins. Clusters were identified on the basis of the estimated 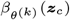 values. (f) Dot plot of enrichment analysis for the top 100 genes upregulated in macrophage Group 2 relative to other groups. (g) UMAP visualization of reconstructed EGFR expression. The red-outlined regions indicate cell populations specifically expressing EGFR. (h) UMAP visualizations of 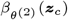 and 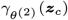. The regions corresponding to the red circles show hazard coefficients that take larger values through synergistic interaction with the BRAF mutation.

Leveraging scTREND’s strength in annotation-free analysis, we conducted an analysis focusing on tumor cells. First, we partitioned cells into two clusters based on 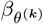 (***z***_*c*_) and identified the cluster with the higher average coefficient. This high-risk-associated cluster was then further subdivided into five clusters, where the number of clusters was determined by maximizing the silhouette score [16] (Figure 3c). Notably, this analysis identified prognosis-associated subclusters based solely on estimated risk coefficients without relying on predefined cell-type categories. Among these sub-clusters, Group 4 exhibited consistently higher risk scores during the early and intermediate time bins compared with the other groups. We therefore performed enrichment analyses on the top 100 genes that were specifically upregulated in Group 4 relative to the remaining tumor cell clusters (Figure 3d). Enrichment analyses for the other groups are shown in Supplementary Figures S2a-d. Group 4 showed enrichment of melanin biosynthesis and MITF-M-dependent gene programs. This finding is consistent with previous reports that MITF-high cells retain proliferative and differentiation capacity [17], demonstrating that scTREND can identify biologically meaningful cell subpopulations in an annotation-free manner.

To validate scTREND’s time-resolved analysis capability, we next focused on macrophages. We applied the same clustering procedure as used for tumor cells. The high-risk cluster was identified by silhouette-based clustering with two clusters (Figure 3e). We then performed functional enrichment analysis on the top 100 genes specifically expressed in Group 2, which exhibited elevated hazard values in time bins 1 and 2 (Figure 3f). This analysis revealed significant enrichment of immune- and inflammation-related pathways, including regulation of immune system processes, inflammatory responses, defense responses, and regulation of cell activation. Macrophages constitute a major immune component of the melanoma tumor microenvironment and are known to contribute to melanoma progression while exerting immunoregulatory functions [18]. While conventional Cox proportional hazards models would have difficulty identifying macrophage subpopulations that exhibit different risk contributions across time intervals, scTREND’s time-resolved approach enabled the capture of immune cell populations that particularly influence prognosis during early stages of disease.

Finally, to demonstrate the utility of scTREND’s condition-dependent modeling, we reconstructed EGFR expression on the latent space using the VAE (Figure 3g) and performed comparative analysis of the patient-shared coefficients 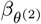(***z***_*c*_) and the BRAF-mutation-specific coefficients 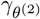 (***z***_*c*_) for the second time interval, and highlighted with red boxes the cell populations exhibiting high reconstructed EGFR expression (Figure 3h). Interestingly, these EGFR-high cell populations showed more pronounced elevated values in the BRAF-mutation-specific coefficient 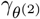 (***z***_*c*_) than in the patient-shared coefficient 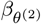 (***z***_*c*_). This is consistent with reports that BRAF inhibitor treatment in BRAF-mutant melanoma can induce activation of the EGFR–SFK–STAT3 signaling axis, contributing to intrinsic or acquired resistance [19]. This pathway acts synergistically with the MAPK cascade, thereby enhancing tumor proliferation and invasion and ultimately worsening patient prognosis. scTREND enables quantitative identification of cell populations whose prognostic risks are amplified through interaction with driver mutations, contributing to the elucidation of mutation-specific risk factors that were previously overlooked by conventional methods.

In summary, scTREND demonstrated its effectiveness in the melanoma cohort from three perspectives: (1) annotation-free identification of prognosis-associated cell subpopulations, (2) detection of cell populations exhibiting different risk contributions across time intervals, and (3) identification of cell populations whose risks are amplified under patient-specific conditions such as BRAF mutations. These results indicate that scTREND possesses high effectiveness and biological interpretability in modeling dynamic and condition-dependent prognostic processes in cancer.

### Analysis of remdesivir-dependent temporal hazard mechanisms in COVID-19 using scTREND

We applied scTREND to bulk transcriptomic data obtained from hospitalized patients with COVID-19. Specifically, bulk transcriptomic data derived from peripheral blood mononuclear cells (PBMCs) were obtained from the IMPACC cohort [20], and single-cell transcriptomic data of PBMCs from [21] were used as the reference scRNA-seq dataset. The cell type annotations from the original study were used (Figure 4a). COVID-19 is an acute respiratory disease caused by infection with the novel coronavirus SARS-CoV-2, and its risk of severe progression is known to vary substantially depending on therapeutic interventions and the time elapsed since disease onset. Such time-dependent and treatment-dependent risk variations in acute diseases provided an ideal case study for validating the effectiveness of scTREND’s time-resolved and condition-aware modeling. In this study, we considered the administration of the antiviral agent remdesivir, one of the therapeutic interventions for COVID-19, as a sample-specific condition. Overall, 59.4% of the patients had received remdesivir treatment. By estimating time- and condition-dependent changes in hazard using our proposed framework, we aimed to gain further insight into the dynamics of disease progression and prognostic factors in COVID-19.

**Fig. 4.**
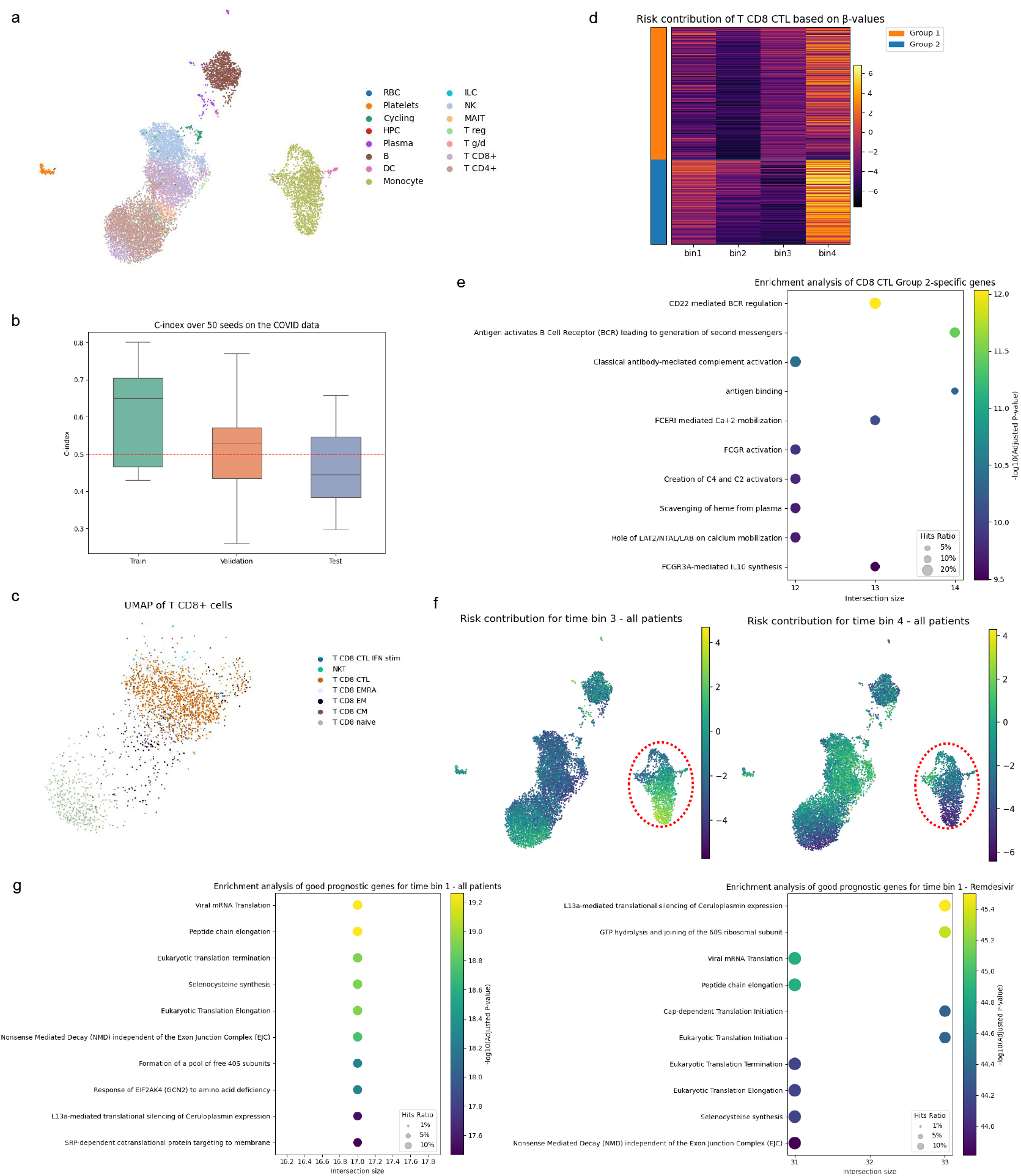
Remdesivir-associated analysis in COVID-19 using scTREND. (a) UMAP visualization of cell types in the reference scRNA-seq dataset. (b) Boxplots of c-index values for the training, validation, and test datasets of the bulk data across 50 random splits. (c) UMAP visualization of refined cell types within CD8^+^ cells. (d) Heatmap of 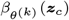 values for CD8^+^ cytotoxic T lymphocytes across time bins. Clusters were identified on the basis of the estimated 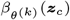 values. (e) Dot plot of enrichment analysis for the top 100 genes upregulated in CD8^+^ cytotoxic T lymphocyte Group 2 relative to other groups. (f) UMAP visualizations of 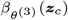 and 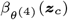. The left panel represents intermediate-phase risk, while the right panel corresponds to late-phase risk. The regions outlined in red indicate monocyte, which show a decrease in hazard coefficients toward the late phase. (g) Dot plots of the top 100 gene set enrichment results negatively correlated with 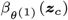 (left) and 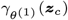 (right) within monocytes.

Across different random splits of bulk samples into training, validation, and test sets, the resulting c-index ranged approximately from 0.4 to 0.7 (Figure 4b). For downstream biological interpretation, we focused on one representative run in which the c-index values on the training and validation sets both exceeded 0.6. The UMAP visualizations of the estimated 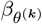 (***z***_*c*_) and 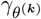 (***z***_*c*_) values at time bins 1-4 are provided in Supplementary Figures S3b and S3c, respectively.

To validate scTREND’s time-resolved analysis capability, we first focused our analysis on CD8^+^ T cells, which play a central role in antiviral immunity. Among them, we further examined cytotoxic T lymphocytes (CTLs), the principal effector population (Figure 4c). Based on the values of 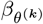 (***z***_*c*_), the CTLs were partitioned into two clusters by maximizing the silhouette score. Notably, through this procedure, scTREND identified a CTL subpopulation that was associated with favorable prognosis during the early and intermediate phases, but caused a pronounced negative effect on prognosis in the late phase (Figure 4d). Such detection of cell populations whose prognostic impact reverses across time intervals is fundamentally impossible with conventional Cox proportional hazards models that assume time-invariance of covariate effects, demonstrating a unique strength of scTREND’s time-resolved approach. To characterize this population, we performed enrichment analysis on the top 100 genes specifically upregulated in this subgroup relative to other CD8^+^ CTLs (Figure 4e). In the late stage of severe COVID-19, immune complexes have been reported to interact with Fc receptors and complement components, contributing to excessive T-cell cytotoxicity and tissue damage [22]. Consistent with this mechanism, enrichment of Fc receptor–related signaling pathways and complement activation was observed in this CTL subgroup. This result demonstrates that the time-dependent risk cell populations statistically identified by scTREND show high biological concordance with known disease mechanisms.

Leveraging scTREND’s strength in annotation-free analysis, we next examined temporal changes in risk contributions across cell types. We visualized the estimated patient shared coefficients 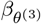 (***z***_*c*_) and 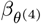 (***z***_*c*_), shared across all bulk samples (Figure 4f). The details of the time point settings are provided in the Methods section. Among these, monocytes exhibited the highest hazard coefficients, suggesting that an increased abundance of monocytes is associated with poor prognosis in COVID-19. This observation is consistent with previous reports showing that elevated levels of monocyte-derived cytokines, such as IL-6 and IL-10, correlate with disease severity in COVID-19 patients [23, 24]. scTREND enabled objective identification of such prognosis-associated cell populations by estimating risk contributions directly from latent representations without relying on predefined cell types.

Finally, to demonstrate the utility of scTREND’s condition-dependent modeling, we focused on monocytes in time bin 1, separately extracting the top 100 genes negatively correlated with the remdesivir-specific hazard coefficient 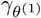 and the patient-shared coefficient 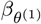, followed by enrichment analyses (Figure 4g). In both the analyses of the entire cohort and the subgroup of patients treated with remdesivir, pathways related to translation (ribosomal processes) and mRNA surveillance were among the most significantly enriched. Previous studies have reported that translational and ribosome-associated pathways, including those involved in translation initiation, tend to be downregulated in severe COVID-19, while being relatively preserved in mild cases [25]. Accordingly, our finding that the maintenance of translational programs in monocytes at early time points is associated with a favorable prognosis is consistent with these observations. Interestingly, stronger enrichment of translation-related pathways was observed in the remdesivir-treated subgroup. Remdesivir treatment has been reported to reduce interferon responses, particularly IFN-*γ*, in hospitalized COVID-19 patients [26], and this finding may reflect treatment-associated modulation of immune and stress-response programs. scTREND’s condition-dependent modeling enabled the identification of prognosis-associated pathways that change specifically with therapeutic intervention, separated from effects common to all patients.

In summary, scTREND demonstrated its effectiveness in the COVID-19 cohort from three perspectives: (1) identification of CTL subpopulations whose prognostic impact reverses across time intervals, (2) objective detection of prognosis-associated cell types in an annotation-free manner, and (3) identification of molecular pathways that change specifically under patient-specific conditions such as remdesivir treatment. In particular, the fact that scTREND functioned effectively in acute infectious disease, a disease domain different from cancer, demonstrates the broad applicability of this framework. These analyses demonstrate that scTREND captures, at single-cell resolution, prognostic contributions that vary with patient-specific conditions and temporal changes, and provides biologically interpretable insights across patients with different treatment histories or clinical backgrounds.

### Dynamic identification of prognosis-associated spatial regions in renal cell carcinoma

We applied scTREND to spatial transcriptomics data from clear cell renal cell carcinoma (ccRCC). Because the tumor immune microenvironment in ccRCC undergoes remodeling during disease progression, we sought to identify spatially defined regions that dynamically contribute to patient prognosis [27]. Such simultaneous analysis of spatial heterogeneity and time-dependent prognostic contributions is difficult to achieve with conventional survival analysis methods, and this provided an ideal case study for validating scTREND’s extensibility to spatial transcriptomics.

We used 10x Visium spatial transcriptomics data and a single-cell RNA-seq dataset from [28], together with bulk RNA-seq data from TCGA for renal cell carcinoma. The cell type annotations from the original study were used (Figure 5a). After predefining four time intervals, consistently high c-index values were obtained across the training, validation, and test sets (Figure 5b). This result demonstrates that scTREND achieves stable predictive performance not only with bulk RNA-seq data but also with spatial transcriptomics data.

**Fig. 5.**
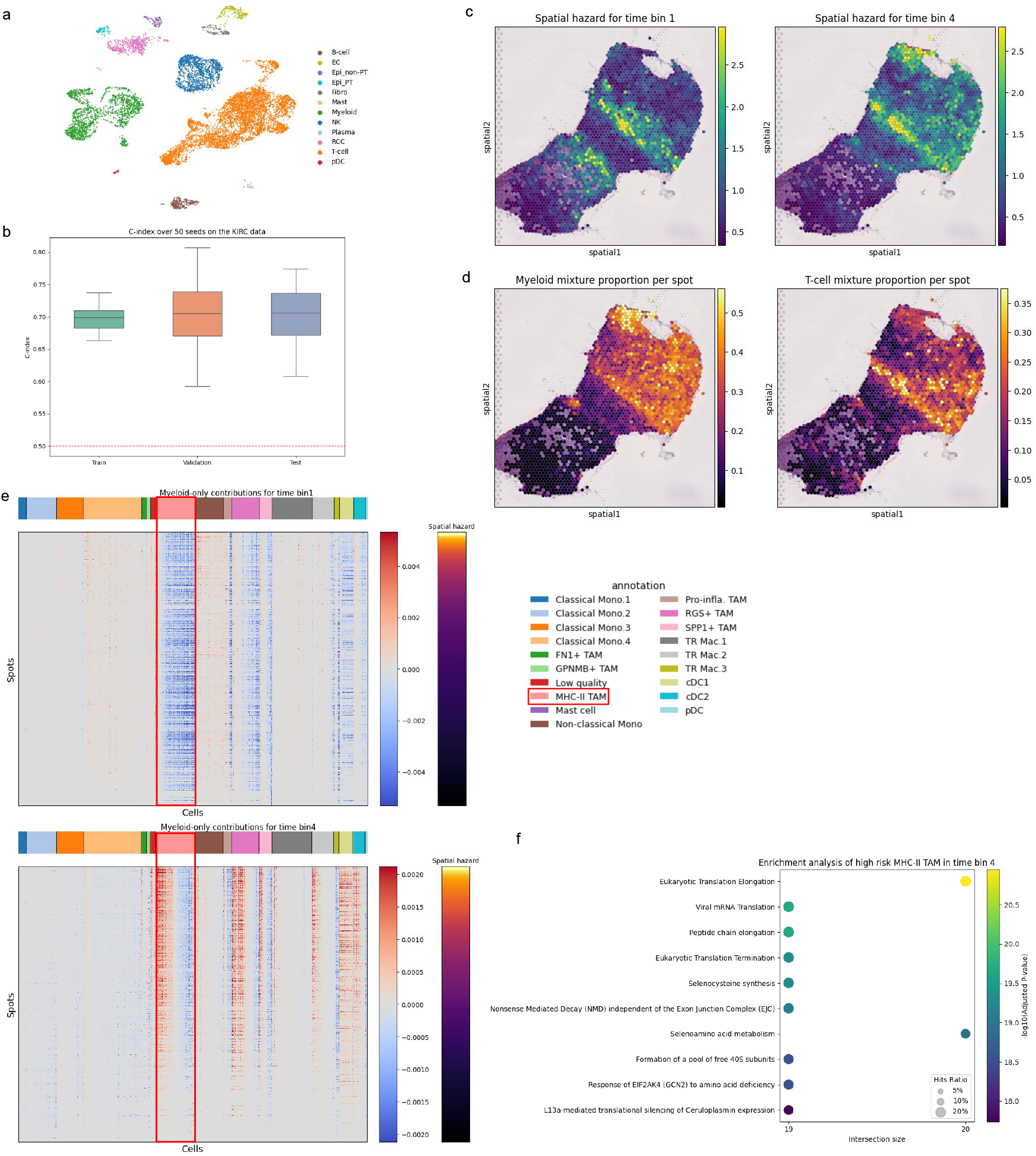
Dynamic spatial analysis of renal cell carcinoma using scTREND. (a) UMAP visualization of cell types in the reference scRNA-seq dataset. (b) Boxplots of c-index values for the training, validation, and test datasets of the bulk data across 50 random splits. (c) Heatmaps of spatial hazard for each spots at time bins 1 (left) and 4 (right). (d) Heatmaps of the spatial proportions of myeloid cells (left) and T cells (right) across spots. (e) Heatmaps of hazard coefficients for each myeloid cell type across spots (time bin 1, upper; time bin 4, lower). (f) Dot plot of enrichment analysis for the top 100 genes upregulated in MHC-II TAM cluster associated with high risk at time bin 4. Clusters were defined by clustering cells based on 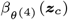 values into high- and low-average-hazard groups.

To demonstrate scTREND’s spatial hazard estimation capability, we calculated spatial hazard values for each spot on the Visium slide. We focused on spatial hazard patterns at time bins 1 and 4 (Figure 5c) and subsequently examined the spatial distributions of Myeloid and T cells across Visium spots (Figure 5d). Spatial hazard estimates for the other time bins and single-cell-level hazard coefficients are provided in Supplementary Figures S4b and S4c, respectively. It is known that, in ccRCC, Myeloid cells exhibit a progression-associated shift from pro-inflammatory to immunosuppressive phenotypes [27]. In parallel, T cells progressively transition toward an exhausted state during disease progression, and both processes have been closely linked to poor clinical outcomes. Notably, scTREND’s analysis revealed that spots exhibiting high spatial hazard values in the late time bin substantially overlapped with regions enriched for both Myeloid and T cells. This result demonstrates that scTREND can effectively integrate spatial cellular composition information with time-dependent risk estimation to identify spatial hazard patterns that are concordant with known biological mechanisms.

Next, focusing on the Myeloid compartment, we examined cells that strongly contributed to the increase in spot-level hazard from the early to the late time point (Figure 5e). This analysis revealed that a subset of tumor-infiltrating macrophages with high MHC class II expression became progressively more risk-associated over time. Tracking the spatial distribution of cell populations whose risk contributions change across time intervals is a unique capability of scTREND that could not be achieved with conventional spatial transcriptomics analysis or survival analysis. To characterize this cell population, we performed enrichment analysis focusing on the top 100 genes that were more specifically expressed compared with other MHC-II-positive tumor-associated macrophages (Figure 5f). In addition to translation-related pathways, including translational elongation, pathways involved in mRNA surveillance and the GCN2-mediated response to amino acid deficiency were significantly enriched. In ccRCC, metabolic reprogramming is known to impose nutrient limitations within the tumor microenvironment [29], and tumor-associated macrophages have been implicated in angiogenesis and immune evasion [30], with increased infiltration and M2-like polarization being associated with poor prognosis [31]. Taken together, the concurrent enrichment of translation-related gene programs and GCN2-mediated nutrient stress responses in this cluster may reflect a TAM state that adapts to nutrient-restricted tumor microenvironments while maintaining tumor-promoting functions. scTREND enabled the identification of such biologically interpretable cell populations by integrating spatial and temporal information.

In summary, scTREND demonstrated its effectiveness in spatial transcriptomics data from ccRCC from three perspectives: (1) identification of spatial hazard patterns that differ across time intervals, (2) detection of spatial regions and cell populations whose prognostic contributions change over time, and (3) identification of biologically interpretable gene programs in spatial and temporal contexts. In particular, the fact that scTREND functioned effectively in spatial transcriptomics data, following the analyses of melanoma and COVID-19 using bulk RNA-seq, demonstrates that this framework possesses versatility that transcends data modalities. These results demonstrate the utility and interpretability of scTREND in linking spatially resolved cellular compositions to dynamic prognostic processes in cancer.

## Conclusion

We introduced scTREND, a deep generative framework designed to estimate time- and condition-dependent risks at single-cell resolution. scTREND integrates a variational autoencoder for latent cellular representation learning, an annotation-free bulk RNA-seq and spatial transcriptome deconvolution module based on DeepCOLOR, and a conditional piecewise constant hazard model that accounts for sample-specific factors.

scTREND possesses three decisive advantages that overcome fundamental limitations of conventional survival analysis methods. First, it can estimate risk contributions at the individual cell level without requiring prior cell-type annotations. Conventional deconvolution methods rely on predefined cell types, potentially missing fine-grained heterogeneity within cell types or prognosis-associated subclusters. scTREND overcomes this constraint by leveraging latent representation learning through VAE. In the melanoma analysis, we successfully identified high-risk tumor cell subclusters expressing MITF-dependent gene programs in an annotation-free manner without relying on predefined cell-type categories. Second, it can dynamically model risk changes over time. Conventional methods such as the Cox proportional hazards model and DeepSurv assume that covariate effects remain constant over time, making them unable to capture changes in prognostic factors accompanying disease progression. The conditional piecewise-constant hazard model in scTREND relaxes this time-invariance assumption. In the COVID-19 analysis, we identified a CTL subpopulation that was associated with favorable prognosis during early and intermediate phases but exhibited negative effects in the late phase. Such detection of cell populations whose prognostic impact reverses across time intervals was fundamentally impossible with conventional methods. Third, it enables risk assessment according to patient-specific conditions such as driver gene mutations and therapeutic interventions. In the melanoma analysis, we successfully identified EGFR-high cell populations whose risks are amplified through interaction with BRAF mutations, and in the COVID-19 analysis, we identified translation-related pathways that change specifically with remdesivir treatment, each separated from effects common to all patients.

Furthermore, by extending this framework to spatial transcriptomics data, scTREND enables the dynamic identification of spatial regions whose prognostic contributions change over time. In the renal cell carcinoma analysis, we revealed that spots exhibiting high spatial hazard values in the late time bin overlapped with regions enriched for Myeloid and T cells, and identified a subset of MHC-II-high tumor-associated macrophages whose risk contributions increase over time. The capability to track the spatial distribution of cell populations whose risk contributions change across time intervals is a unique strength of scTREND that could not be achieved with conventional spatial transcriptomics analysis or survival analysis.

In numerical simulations, scTREND accurately recovered both time-dependent and condition-dependent hazard coefficients, achieving predictive performance superior to the existing method scSurv. In applications to real datasets, the identified cell populations and molecular pathways showed high concordance with known biological mechanisms, including the EGFR–SFK–STAT3 signaling pathway, Fc receptor-related signaling, and GCN2-mediated nutrient stress responses. Of particular note, scTREND functioned effectively and consistently across different disease domains—cancer (melanoma, renal cell carcinoma) and acute infectious disease (COVID-19)—as well as across different data modalities—bulk RNA-seq and spatial transcriptomics—demonstrating the broad versatility of this framework.

Despite these advantages, several challenges remain. The stability of hazard estimation may depend on the fidelity of bulk deconvolution and on the number of observed events within each time interval. Furthermore, the present implementation considers a single conditioning factor, whereas future extensions should enable modeling of multiple interacting conditions, such as the combined effects of genetic mutations and therapeutic interventions.

In summary, scTREND provides an innovative framework for single-cell survival analysis that combines three key features: annotation-free, time-resolved, and condition-aware analysis. By linking single-cell states to clinical outcomes along the temporal axis, scTREND can identify cell populations and pathways associated with prognosis, thereby contributing to the development of therapeutic strategies tailored to disease stage and progression.

## Methods

### scTREND architecture

The scTREND architecture comprises three modules: (i) latent representation learning from scRNA-seq using a variational autoencoder (VAE), (ii) estimation of sample-specific cellular composition via bulk RNA-seq deconvolution based on DeepCOLOR [32], and (iii) risk modeling using a conditional piecewise-constant hazard model [33] to capture condition-dependent and time-dependent effects. Together, these modules provide an end-to-end pipeline that transforms single-cell-derived latent states into condition-dependent and time-resolved patient-level risk predictions.

Specifically, the VAE encodes scRNA-seq profiles into a low-dimensional latent space representing biological cellular states. DeepCOLOR then maps bulk RNA-seq samples to cell assignment probabilities, which serve as composition weights. Finally, the conditional piecewise-constant hazard model computes patient-level risk scores by integrating these composition weights with the latent representations.

For spatial transcriptomics, the deconvolution module is replaced by cell assignment to spatial spots. The resulting spot-wise composition weights are used in the same survival model to estimate spatially resolved hazards and to identify prognosis-associated regions across time intervals. The following subsections describe these components sequentially.

#### Variational Representation of Single cell

We begin by considering a generative model for scRNA-seq expression data. We denote the scRNA-seq expression matrix as ***X***∈ ℝ^*C*×*G*^. Let *C* and *G* denote the number of cells and genes, respectively. Assuming there are *P* batches (e.g., patients), we represent the batch information for cell *c* as ***p***_*c*_ [0, 1]^*P*^. Because scRNA-seq expression data are inherently high-dimensional, directly using them for hazard estimation can lead to overfit and poor generalization. Therefore, we employ the conditional VAE to obtain a low-dimensional latent variable ***z***_*c*_ ∈ℝ^*d*^ for each cell *c* [34]. We assume that the observed gene expression count *x*_*cg*_ for gene *g* in cell *c*, given the latent representation ***z***_*c*_ and the batch information ***p***_*c*_, is distributed according to a Poisson distribution defined as follows:

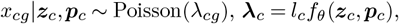

where *l*_*c*_ ∈ ℝ represents the library size, i.e., the mean expression across all genes in cell *c*, and *f*_*θ*_ : ℝ*d*+^*P*^ → ℝ^*G*^ is implemented as a multi-layer perceptron serving as the decoder, translating the latent cell state back into gene space. As the prior distribution, each component of the latent variable ***z***_*c*_ is assumed to be drawn from a mutually independent standard normal distribution:

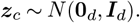

The posterior distribution of ***z***_*c*_ is parameterized by an encoder neural network that outputs the mean and variance, denoted as *μ*_*ϕ*_(***x***_*c*_, ***p***_*c*_) and *σ*_*ϕ*_(***x***_*c*_, ***p***_*c*_), respectively:

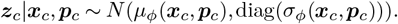

Based on these formulations, the evidence lower bound (ELBO) is defined as follows:

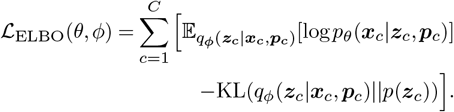

The expectation term is approximated using the reparameterization trick, where a single Monte Carlo sample per mini-batch is used during training.

#### Annotation-free bulk and spatial deconvolution

Now, we discuss the method of the deconvolution for bulk RNA-seq data. Let *B* denote the number of samples in the bulk RNA-seq dataset. Given the latent representation of each cell ***z***_*c*_, the cellular composition at the single-cell level for each bulk sample is estimated using DeepCOLOR. In DeepCOLOR, the observed gene expression value *e*_*bg*_ for gene *g* in bulk sample *b* is assumed to follow a negative binomial distribution as follows:

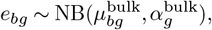

where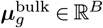denotes the mean parameter, which is parameterized as

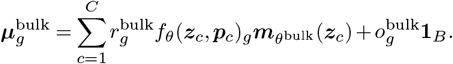

***r***^bulk^∈ℝ^*G*^ represents gene-specific correction factors adjusting for capture efficiency differences between scRNA-seq and bulk RNA-seq data. The mapping 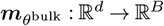is implemented as a neural network that takes the latent representation of a cell as input and outputs its assignment probabilities across bulk samples. Accordingly, for all bulk samples *b*, the normalization constraint 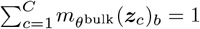 holds. In addition, ***o***^bulk^ ∈ ℝ ^*G*^ denotes gene-specific shift parameters, and ***α***^bulk^ ∈ ℝ^*G*^ represents dispersion parameters. Based on these definitions, the negative log-likelihood (NLL) is formulated as

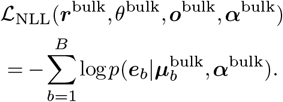

Next, we describe the spatial deconvolution procedure. Let *S* denote the number of Visium spots in the spatial transcriptome data. Similar to the bulk case, the observed gene expression value *e*_*sg*_ for gene *g* in spot *s* is assumed to follow a negative binomial distribution as follows:

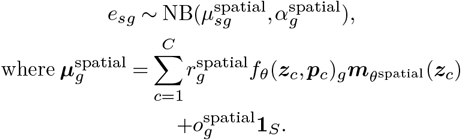

All model parameters are defined in the same manner as in the bulk setting. In particular, the mixture proportions for each spot *s* are constrained to sum to one, i.e., 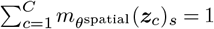.The NLL is formulated as

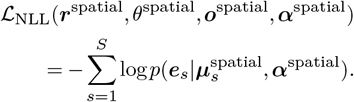

In the spatial transcriptomics analysis, we defined the spatial hazard for each spot on the Visium slide as a softplus-transformed product of the estimated single-cell mixture proportions and the corresponding hazard coefficients.

#### Single-cell time- and condition-dependent hazard modeling

Using the latent representations ***z***_*c*_ obtained from the VAE and the cellular mixing proportions ***m***_*θ*bulk_(***z***_*c*_) estimated by DeepCOLOR, we estimate the time-dependent, single-cell level hazard while accounting for sample-specific factors such as driver gene mutations or treatment history. This model can be regarded as a neural network-based piecewise constant hazard model that is conditioned on sample-specific factors. We define discretized time intervals as *t*_0_(= 0) *< t*_1_ *<… < t*_*K*_ with Δ_*k*_ = *t*_*k*_*−t*_*k−*1_, which must be pre-specified by the analyst. For each bulk sample *b*, let *T*_*b*_ denote the observed time and *δ*_*b*_ the event indicator. The corresponding time interval index is defined as {*k*|(*T*_*b*_) = min *k T*_*b*_ ≤ *t*_*k*_}. To parameterize both the patient-shared effect and the condition-specific hazard associated with sample-specific factors in interval *k*, we define the linear predictor *η*_*bk*_ as follows:

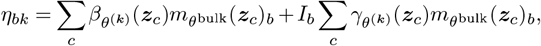

where *I*_*b*_ is an indicator function that returns 1 if bulk sample *b* satisfies a sample-specific condition, and 0 otherwise. The hazard rate for bulk sample *b* in interval *k* is then defined as *λ*_*bk*_ = softplus(*η*_*bk*_), such that the hazard function can be expressed as *h*_*b*_(*t*) = *λ*_*bk*_. The cumulative hazard Λ_*b*_(*t*) and survival function *S*_*b*_(*t*) are defined as

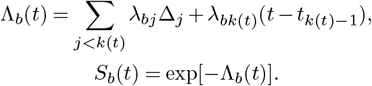

Accordingly, the probability density function of the event time for sample *b* at time *t* is *f*_*b*_(*t*) = *λ*_*bk*_(*t*)*S*_*b*_(*t*). Based on these definitions, the NLL is formulated as

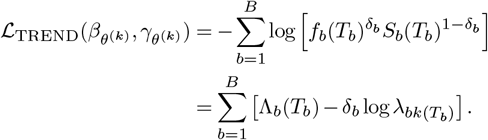

### Training procedure and optimization strategy

Model training was performed in three stages using the AdamW optimizer, each corresponding to the objectives described above. Early stopping was applied in all stages: training was terminated when the moving average of the validation loss over the most recent 30 epochs failed to improve for 30 consecutive evaluations. In the first stage, which involved training the VAE, the mini-batch size was set to 1,000, the learning rate to 0.01. The encoder network comprised a hidden layer of 100 units and a latent space of 20 dimensions. The scRNA-seq dataset was divided into training, validation, and test sets at a ratio of 0.85:0.10:0.05. In the second stage, corresponding to DeepCOLOR training, the mini-batch size and learning rate were again set to 1,000 and 0.01. In the third stage, which estimated the hazard coefficients, the bulk RNA-seq datasets were split into training, validation, and test sets at a ratio of 0.6:0.2:0.2, with a mini-batch size of 500 and a learning rate of 0.0001. To prevent the cumulative hazard from diverging due to excessively wide time intervals, survival times were first trimmed by temporarily excluding the top and bottom 1% of samples. The remaining samples were then min-max scaled to the range [0, 5], after which the previously excluded samples were reassigned values of 5 (top 1%) and 0 (bottom 1%), respectively. For applications of scTREND to the melanoma, COVID-19, and renal cell carcinoma datasets, the time intervals were set to [0, 0.5, 1.0, 2.0, 5.0], [0, 0.1, 0.2, 0.4, 5.0], and [0, 0.5, 1.0, 2.0, 5.0], respectively, to ensure adequate numbers of events within each interval. The corresponding real-time durations for these normalized intervals are provided in Supplementary Figures S1a, S3a, and S4a.

### Simulation study design and evaluation metrics

#### Pseudo-bulk data generation

Simulated data were generated using the breast cancer scRNA-seq dataset from [10], following the simulation procedure described in scSurv [11]. First, we selected the patient with the largest number of recorded cells and randomly split the corresponding dataset into two subsets. One subset was used as the reference scRNA-seq data, and the other was employed for pseudo-bulk data generation. For pseudo-bulk generation, we used the predefined cell-type labels “celltype_minor” as clusters. A probability vector was sampled from a Dirichlet distribution with hyperparameter *α* = 1. The total number of cells for each pseudo-bulk sample was then drawn from a uniform distribution *U* (1000, 10000). Using the previously sampled probability vector as parameters, cell-type compositions were determined by multinomial sampling. Within each cell type, cells were sampled with replacement and aggregated to form a pseudo-bulk sample. For each pseudo-bulk sample *b*, the proportion of cell type *n* was calculated and denoted as *m*_*bn*_, satisfying 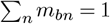.

#### Survival time simulation

True parameter values *β*_*kn*_ and *γ*_*kn*_ for each cell type *n* in time interval *k* were generated through independent random walks. Each parameter was initialized at *β*_0*n*_ and *γ*_0*n*_, respectively, and updated as follows:

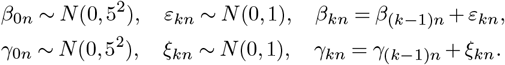

The follow-up period was divided into five time points {0, 0.5, 1.0, 2.0, 5.0}, which define four piecewise-constant hazard intervals: [0, 0.5), [0.5, 1.0), [1.0, 2.0), and [2.0, 5.0). This discretization helps avoid explosive growth of the cumulative hazard when the interval length Δ_*k*_ = *t*_*k*_ *− t*_*k−*1_ becomes large. In practice, the time axis can be rescaled or normalized depending on the dataset, since such transformation does not affect the model or its inference. For each pseudo-bulk sample *b*, we sampled a binary indicator *I*_*b*_ ∈{0, 1}representing a sample-specific condition from a Bernoulli distribution with success probability 0.5. Using the true parameters, we then defined the hazard function *h*_*b*_(*t*) for bulk *b* and its cumulative hazard *H*_*bk*_ up to interval *k* as:

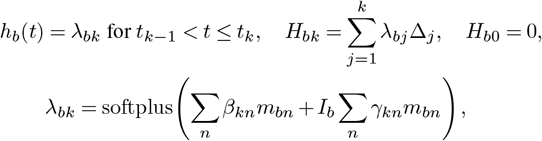

where *m*_*bn*_ denotes the proportion of cell type *n* in bulk *b*, satisfying 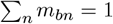. Event times were simulated using the inverse transform sampling method. For each bulk *b*, we first sampled *U*_*b*_ *∼U* (0, 1) and set *Z*_*b*_ = *−*log *U*_*b*_. The event time *T*_*b*_ was then determined by

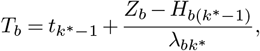

where *k*^*∗*^ = min {*k* : *H*_*bk*_ ≥*Z*_*b*_}. Finally, we treated observations with *T*_*b*_ ≥5 as right-censored (*δ*_*b*_ = 0), and the remaining ones as events (*δ*_*b*_ = 1).

#### Evaluation metrics

Using the pseudo-bulk samples generated in the previous subsection, we varied the true regression coefficients and simulated 50 distinct sets of survival times. To evaluate the estimation performance, we computed the Pearson correlation between the true and estimated regression coefficients, as well as the concordance index (c-index) based on the risk score 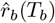 defined as

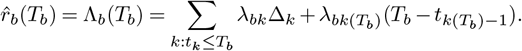

## Author contributions statement

S.Y. conceived the core model underlying the proposed methodology and implemented the source code, as well as conducted performance evaluation, under the supervision of C.M. and T.S. K.A. contributed to the further development and refinement of the methodology. All authors reviewed and approved the final manuscript.

## Acknowledgments

We gratefully acknowledge the IMPACC study group (IMPACC Manuscript Writing Team and IMPACC Network Steering Committee 2021) for providing bulk RNA-seq data and associated clinical outcome data used in this study.

## Data and code availability

The implementation of scTREND is available at https://github.com/R301Carbine/scTREND. The scRNA-seq datasets used for the breast cancer and melanoma analyses can be accessed from GSE176078 and GSE115978, respectively. The bulk RNA-seq data and corresponding clinical information for renal cell carcinoma and melanoma were obtained from the Genomic Data Commons (GDC) Data Portal, whereas the COVID-19 bulk RNA-seq dataset and clinical metadata were retrieved from ImmPort. The preprocessing procedures for all datasets used in this study followed the protocols described in [11]. Specifically, the preprocessing of the breast cancer scRNA-seq dataset from [10], which was used for numerical simulations, and those of the melanoma, COVID-19, and renal cell carcinoma datasets employed in the real-data analyses, were performed according to the subsections “scRNA-seq data preprocessing,” “TCGA bulk data preprocessing,” “COVID-19 PBMC bulk data preprocessing,” and “Spatial transcriptome preprocessing,” respectively, in [11]. Functional enrichment analysis was performed using g:Profiler [35], using Gene Ontology Biological Process terms [36, 37] and Reactome pathways [38].

---

**Fig. S1.**
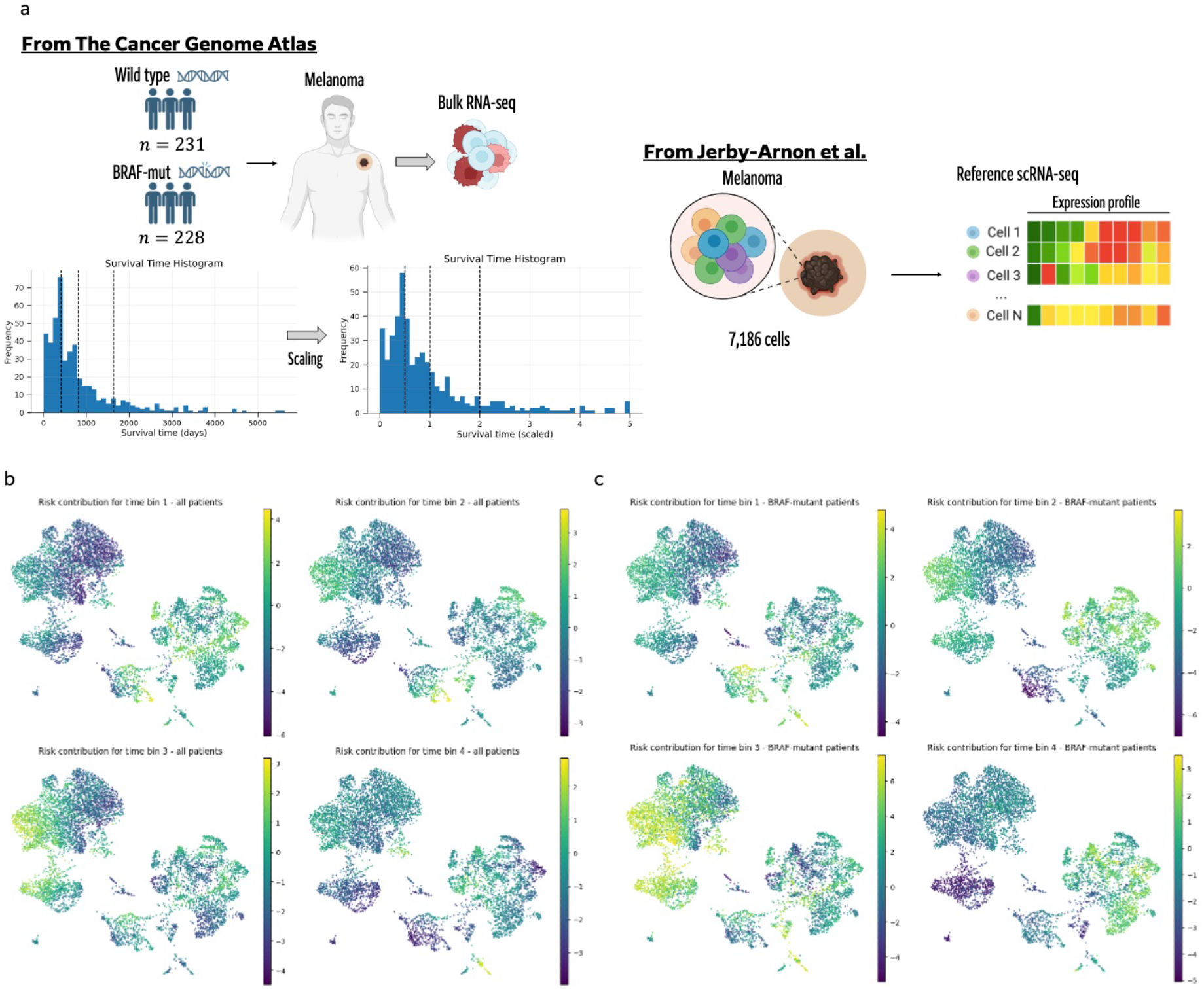
Time-resolved UMAP visualizations of patient-shared and BRAF-mutant-specific coefficients in melanoma dataset. (a)Dataset and time-interval settings for survival analysis. The melanoma scRNA-seq and bulk RNA-seq data were used, conditioned on BRAF mutation status. Time intervals for survival analysis were defined to ensure a sufficient number of subjects in each interval, with the interval boundaries indicated by dashed lines in the histogram. (b) UMAP visualizations of the patient-shared coefficient 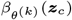 across time bin 1-4 in the melanoma dataset. (c) UMAP visualizations of the BRAF-mutant-specific coefficient 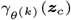 across time bin 1-4 in the melanoma dataset.

**Fig. S2.**
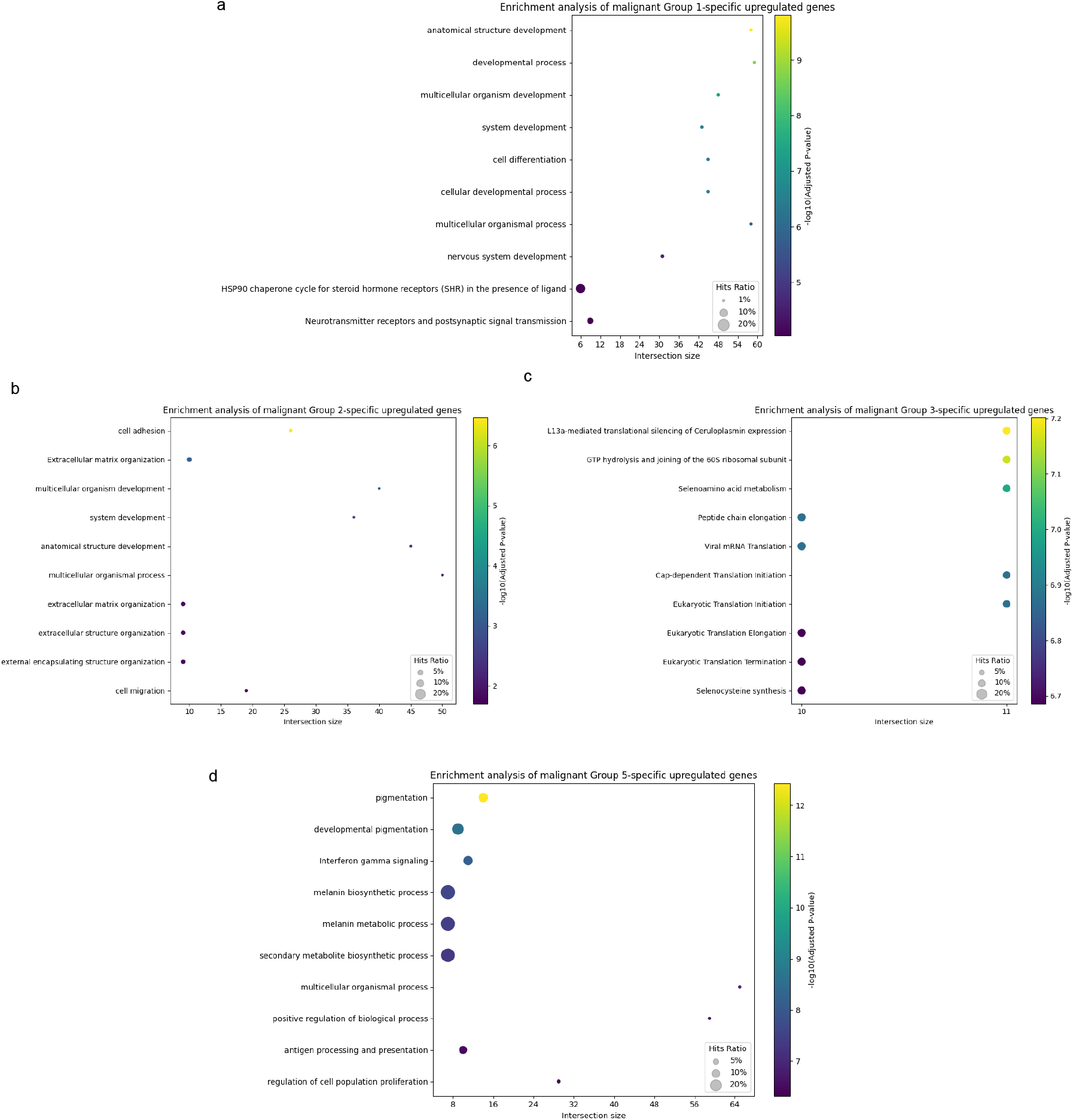
Enrichment analysis of malignant subgroups identified in the melanoma dataset. (a) Dot plot of enrichment analysis for the top 100 genes specifically upregulated in malignant cell Group 1 relative to other groups. Each dot represents an enriched gene set; the y-axis lists enriched terms, and the x-axis indicates the number of overlapping genes. Dot size reflects the ratio of overlapping genes to gene set size, and color intensity represents statistical significance. (b) Dot plot of enrichment analysis for the top 100 genes upregulated in malignant cell Group 2 relative to other groups. (c) Dot plot of enrichment analysis for the top 100 genes upregulated in malignant cell Group 3 relative to other groups. (d) Dot plot of enrichment analysis for the top 100 genes upregulated in malignant cell Group 5 relative to other groups.

**Fig. S3.**
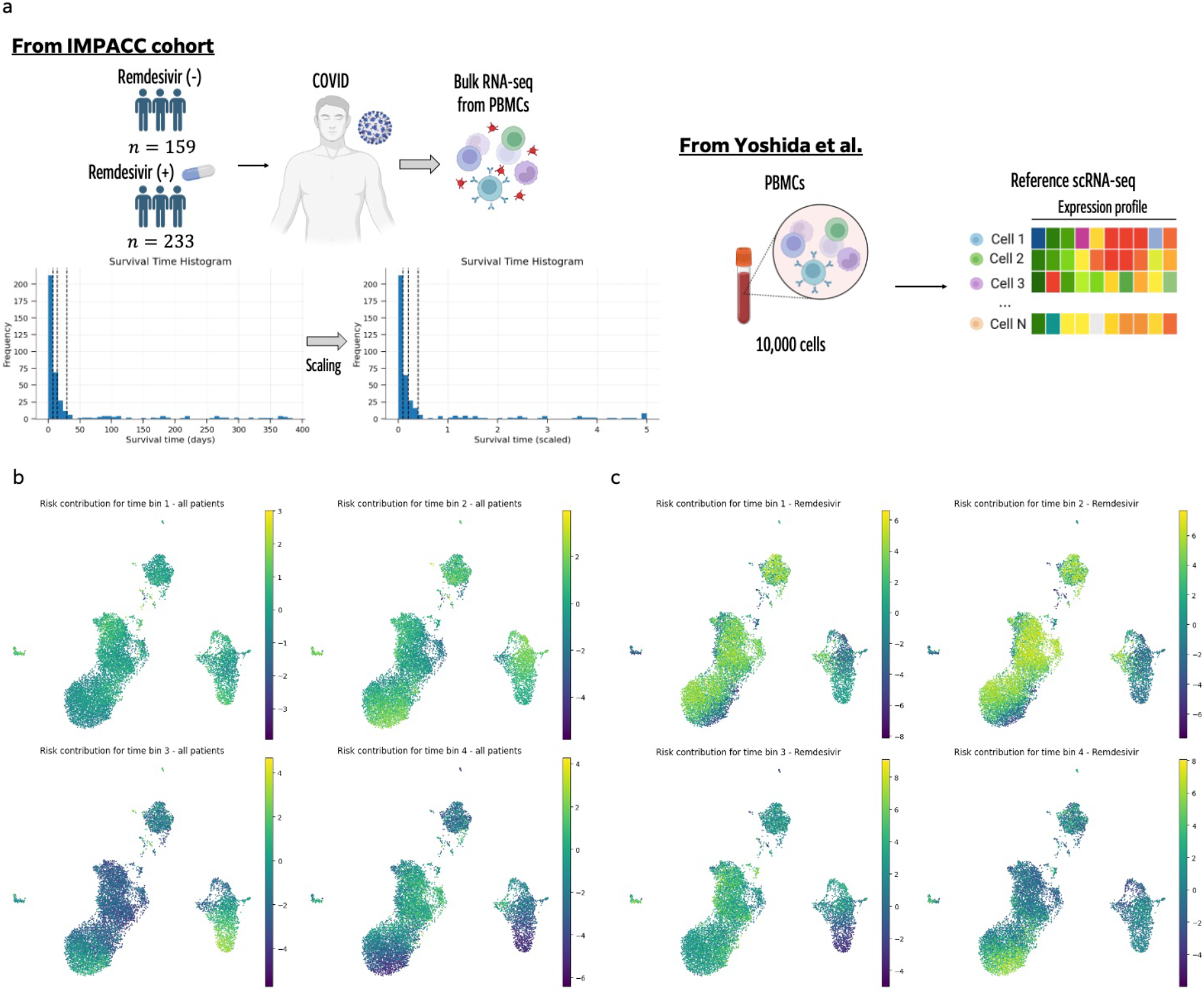
Time-resolved UMAP visualizations of patient-shared and remdesivir-treated-specific coefficients in COVID dataset. (a) Dataset and time-interval settings for survival analysis. The COVID-19 peripheral blood mononuclear cell (PBMC) scRNA-seq and bulk RNA-seq data were used, conditioned on remdesivir treatment. Time intervals for survival analysis were defined to ensure a sufficient number of subjects in each interval, with the interval boundaries indicated by dashed lines in the histogram. (b) UMAP visualizations of the patient-shared coefficient 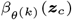 across time bin 1-4 in the COVID dataset. (c) UMAP visualizations of the remdesivir-treated-specific coefficient 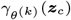 across time bin 1-4 in the COVID dataset.

**Fig. S4.**
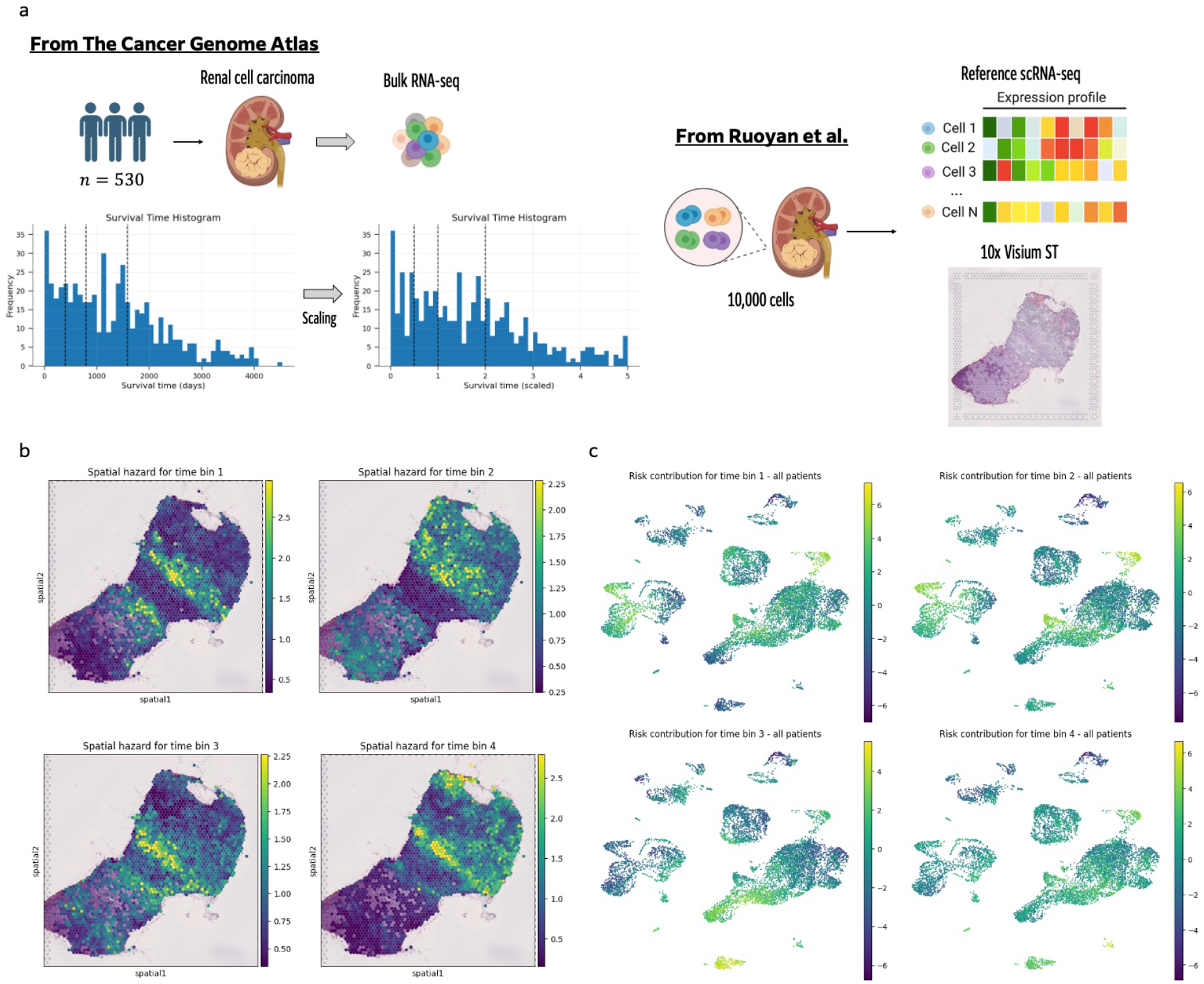
Time-resolved spatial hazard and UMAP visualizations of patient-shared coefficients in the renal cell carcinoma dataset. (a) Dataset and time-interval settings for survival analysis. Renal cell carcinoma scRNA-seq, bulk RNA-seq, and 10x Visium spatial transcriptomics data were used. Time intervals for survival analysis were defined to ensure a sufficient number of subjects in each interval, with the interval boundaries indicated by dashed lines in the histogram. (b) Heatmaps of spatial hazard across time bin 1-4 in the renal cell carcinoma dataset. (c) UMAP visualizations of the patient-shared coefficient 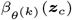 across time bin 1-4 in the renal cell carcinoma dataset.

